# Activity-gated exocytosis drives myelin remodeling required for adult learning

**DOI:** 10.64898/2026.10.19.760195

**Authors:** Mable Lam, Oleh Dehtiarov, Anna C. Geraghty, Belgin Yalçın, Lauren Duan, Manasi Iyer, Kristen H. Schuster, Michelle Monje, Ethan G. Hughes, J. Bradley Zuchero

**Affiliations:** Department of Neurosurgery, Stanford University School of Medicine; Stanford, CA, USA; Department of Neurology and Neurological Sciences, Stanford University School of Medicine, Stanford, CA, USA; Howard Hughes Medical Institute, Stanford, CA, USA; Department of Cell and Developmental Biology, University of Colorado School of Medicine, Aurora, CO, USA

## Abstract

Myelin remodeling is emerging as a dynamic facet of plasticity in the adult brain. Learning and sensory experience induce remodeling of existing myelin sheaths through incompletely understood mechanisms, leaving its causal contribution to adaptive behaviors unclear. Here, we identify activity– dependent exocytosis in mature oligodendrocytes as a key driver of myelin remodeling. Live-cell imaging reveals that neuronal activity stimulates oligodendrocyte exocytosis. Exocytosis in mature oligodendrocytes drives myelin sheath elongation on optogenetically stimulated axons in adult mice. Disrupting exocytosis in pre-existing myelin, but not newly formed myelin, impairs motor skill acquisition, suggesting that remodeling of pre-existing myelin sheaths is required for learning. These results uncover a cellular mechanism that couples neuronal activity to myelin membrane remodeling and provide a molecular handle to interrogate the function of myelin plasticity across different circuits, behaviors, and neurological disorders.

## Introduction

Myelin governs circuit timing and is an integral component of structural plasticity in the brain. Learning and experience can add new myelin through generating new oligodendrocytes^1–3^ and remodel the structure of pre-existing sheaths^4–6^, changes thought to fine-tune the timing and synchrony of action potential conduction in neural circuits^7,8^. Causal evidence links new myelin formation to learning, as blocking oligodendrocyte generation impairs many forms of learning and memory^1,2,9–12^. Whether remodeling of pre-existing sheaths is likewise required is unknown, largely because tools to perturb pre-existing myelin in an activity-dependent manner have been lacking^8^. Importantly, evidence of continuous myelin renewal with only low levels of new oligodendrocyte generation from human brain samples^13^ suggests that myelin remodeling may be an active mechanism of myelin plasticity in humans, underscoring the need to determine its causal role in circuit function and adaptation.

Advancing our understanding of myelin remodeling requires identifying cellular mechanisms that drive activity-dependent changes to myelin morphology of the adult brain. Exocytosis is a well-established, activity-regulated process in many cell types, most notably during synaptic vesicle release at synapses^14,15^. We previously reported that oligodendrocytes require exocytosis to add new membrane during myelin sheath formation in development^16^. Notably, we observed that exocytosis adds new myelin membrane at axon-myelin interfaces that occur along sheath edges and within the innermost myelin wrap, suggesting that the exocytosis machinery may be poised to respond to neuronal activity to promote sheath remodeling.

Here, we combine live-cell imaging, sparse labeling of myelin in mouse brains, and behavioral assays to define the mechanisms of myelin remodeling and its role in learning. We first use chemogenetic tools in live-cell imaging and find that glutamatergic neuronal activity more than doubles the rate of exocytosis in mature oligodendrocytes. Then, through optogenetic stimulation of neurons in mice, we find that exocytosis is required for myelin sheath elongation on activated axons in the corpus callosum. Disrupting exocytosis in mature oligodendrocytes to block myelin remodeling impaired motor skill learning in adult mice, while disrupting exocytosis in precursors (OPCs) and newly formed oligodendrocytes exclusively impaired motor memory without affecting learning. Our findings reveal a key pathway to target remodeling of pre-existing myelin and demonstrate its causal contribution to learning in the adult brain.

## Results

### Glutamatergic neuronal activity stimulates exocytosis in mature oligodendrocytes

To test whether neuronal activity modulates oligodendrocyte exocytosis, we performed live-cell imaging of co-cultures pairing oligodendrocytes with reaggregated rat retinal ganglion cell–derived neurons^17^ expressing hM4Di inhibitory DREADD (designer receptors exclusively activated by designer drugs)^18^ to silence activity with clozapine-N-oxide (CNO) (Fig. 1a). Acute CNO treatment of hM4Di-infected retinal ganglion cells suppressed calcium transients in neurites, as expected (Extended Data Fig. 1a-b). We measured exocytosis in oligodendrocytes by expressing VAMP2-or VAMP3-conjugated pHluorin, the pH-sensitive variant of GFP^14,19^ which reports events of vesicle fusion to the plasma membrane as transient, punctate increases in fluorescence (Fig. 1b). Compared to oligodendrocytes in isolation, we found that co-culturing with neurons increases the average rate of VAMP2 exocytosis by 1.7-fold and of VAMP3 exocytosis by 2.8-fold in oligodendrocytes (Fig. 1c-d). Interestingly, silencing neuronal activity reduced VAMP3, but not VAMP2, exocytosis rates to the levels seen in oligodendrocytes cultured alone.

**Fig. 1.**
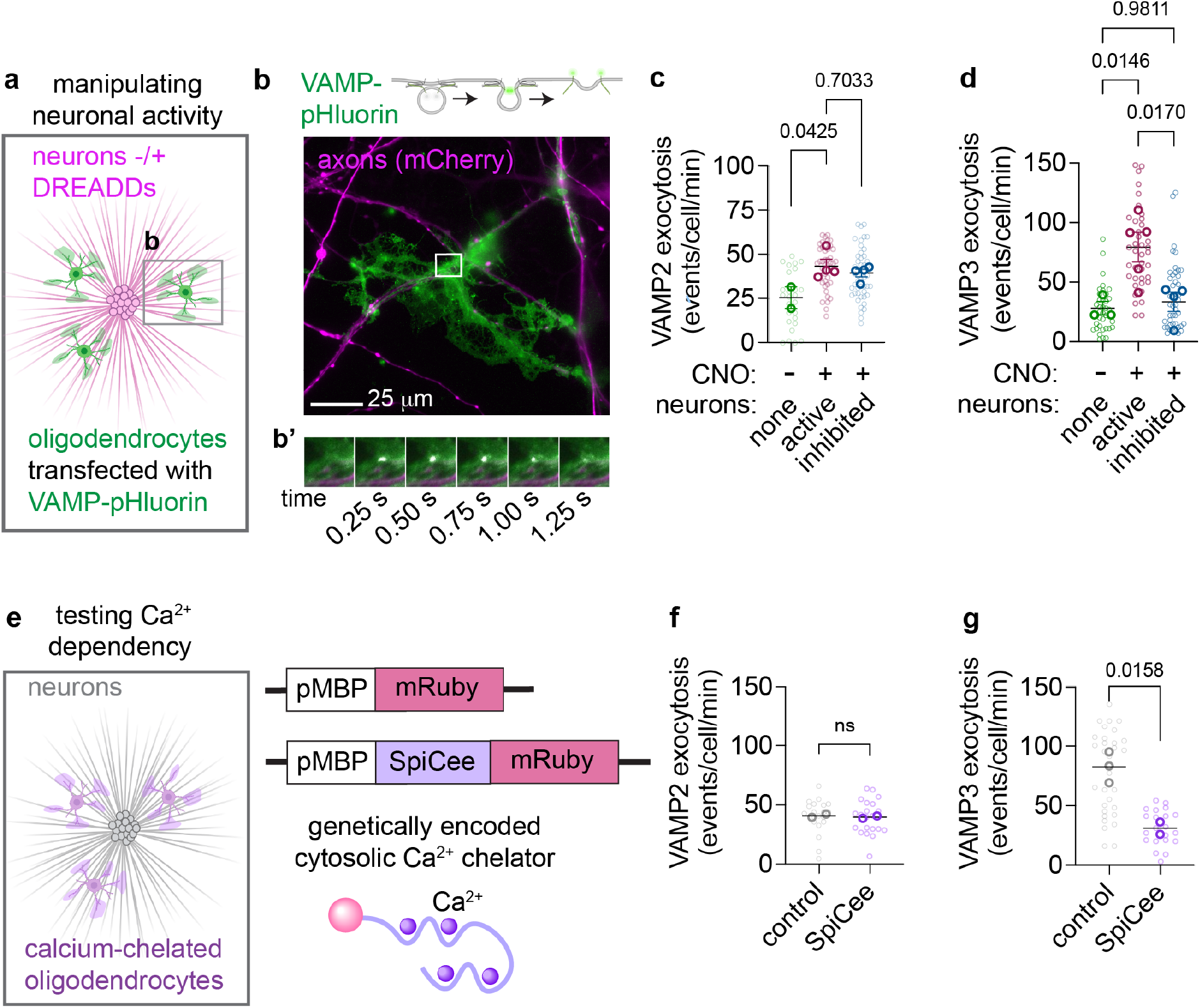
Neuronal activity increases calcium-dependent exocytosis in mature oligodendrocytes. (**a**) Schematic of primary co-cultures between oligodendrocytes (green) expressing the pH-sensitive exocytosis reporter (VAMP-pHluorin) and neurons (magenta) from re-aggregated retinal ganglion cells expressing mCherry (control) or DREADD hM4Di-mCherry to inducibly silence activity. (**b**) A single oligodendrocyte expressing VAMP3-pHluorin (green) ensheathing axons (magenta) at after 5 days of co-culture. Inset (b’) highlights a single exocytotic event marked by a transient punctate increase of fluorescence that decays over time. (**c**,**d**) Exocytosis rates of VAMP2 (c) and VAMP3 (d) in oligodendrocytes cultured for 5 days in isolation (green), with active mCherry-expressing neurons (magenta), or with DREADD hM4Di-expressing neurons (blue). Co-cultures were treated with 50 nM clozapine-N-oxide (CNO) for 1 hr prior to imaging. p-values: one-way ANOVA with Tukey correction for multiple comparisons. (**e**) Schematic of primary co-cultures between uninfected neurons (grey) and oligodendrocytes (purple), which express cytosolic mRuby (control) or the genetically encoded calcium chelator SpiCee conjugated to mRuby. (**f**,**g**) Exocytosis rates of VAMP2 (f) and VAMP3 (g) in oligodendrocytes expressing mRuby (control) or SpiCee (calcium chelator) co-cultured for 5 days with active neurons. p-values: unpaired, two-tailed t-test. See Supplementary Table 1 for average event frequencies ± SEM.

How might neuronal activity regulate VAMP3-mediated exocytosis in oligodendrocytes? Neuronal activity evokes calcium transients in myelin sheaths in vitro and in vivo during development^20–22^, and calcium triggers regulated exocytosis of select vesicle populations in many cell types^15^. To test whether calcium regulates activity-dependent exocytosis in oligodendrocytes, we suppressed calcium transients specifically in oligodendrocytes by expressing the genetically encodable cytosolic calcium chelator SpiCee^23,24^ together with VAMP-conjugated pHluorin (Fig. 1e). Suppressing oligodendrocyte calcium transients in the presence of active neurons significantly reduced VAMP3, but not VAMP2, exocytosis (Fig. 1f-g), mirroring the effect of silencing neuronal activity.

To identify the neuronal signal that stimulates calcium-dependent exocytosis in oligodendrocytes, we first determined if increasing exocytosis requires a secreted factor or an axon membrane-bound factor. We compared the exocytosis rates of oligodendrocytes that contact axons (contacting) to oligodendrocytes that make no contact (non-contacting) within a single imaging well (Fig. 2a). We found that contacting and non-contacting oligodendrocytes exhibited similar rates of VAMP3 exocytosis (Fig. 2b-c), suggesting that a secreted neuronal factor drives activity-dependent exocytosis in oligodendrocytes. Because the co-cultured retinal ganglion cells are glutamatergic and glutamate evokes calcium transients in oligodendrocytes via NMDA (N-methyl-d-aspartate) receptor signaling in rodent slice culture^25,26^ and metabotropic glutamate receptor 5 (mGluR5) in zebrafish^21^, we hypothesized that glutamate stimulates exocytosis in oligodendrocytes. Acute treatment of isolated oligodendrocytes with glutamate increased the exocytosis rate of VAMP3 by 1.8-fold but not of VAMP2 (Fig. 2d). Co-treatment with the mGluR5 antagonist MTEP suppressed this glutamate-induced increase in VAMP3 exocytosis, whereas the NMDA receptor antagonist MK-801 or AMPA receptor antagonist NBQX had little effect (Fig. 2e). Altogether, these results reveal that the presence of neurons enhances both VAMP2 and VAMP3 exocytosis in oligodendrocytes but that VAMP3 is selectively tuned by activity-dependent calcium transients through glutamate receptor signaling.

**Fig. 2.**
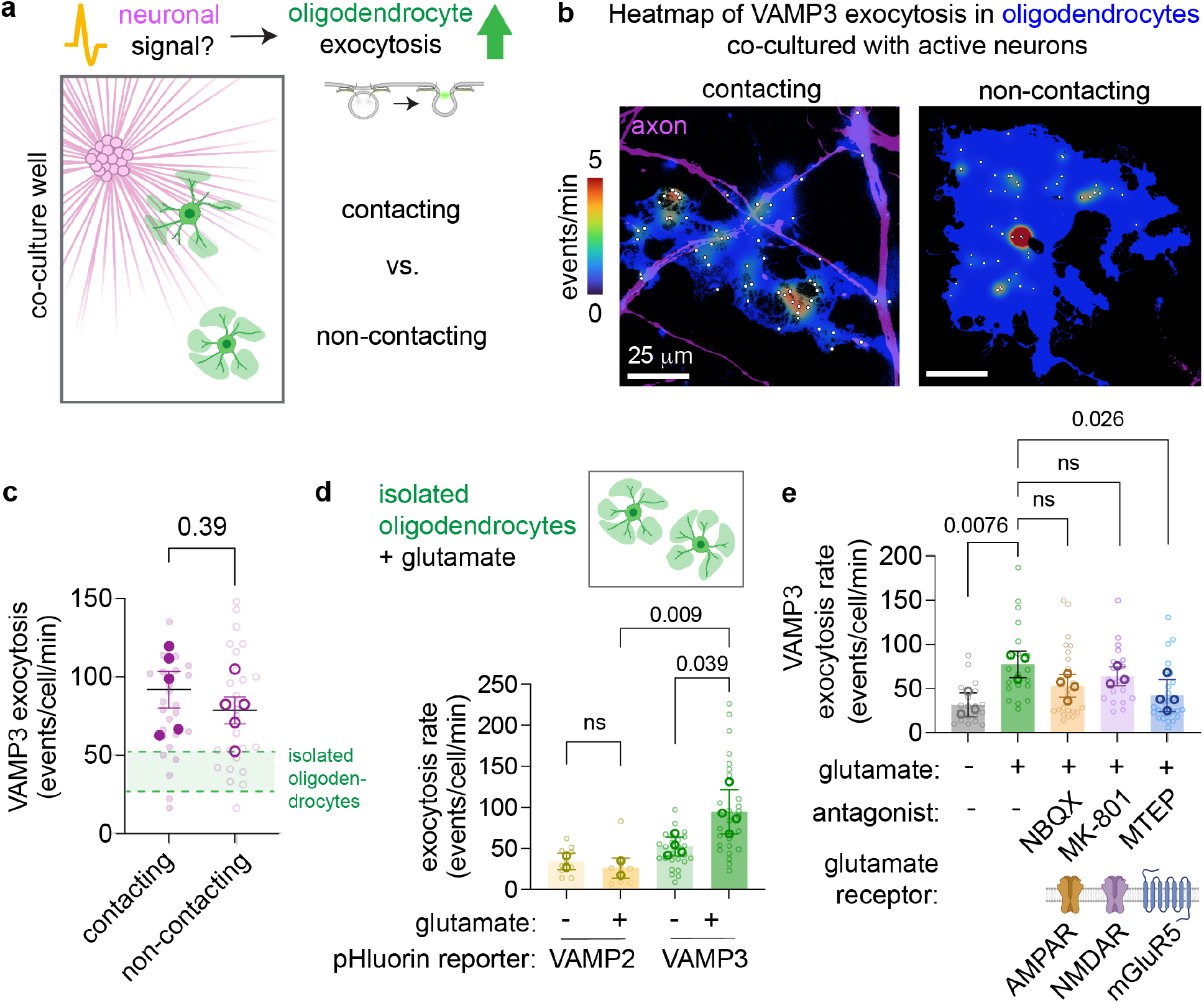
The neurotransmitter glutamate induces activity-dependent exocytosis in oligodendrocytes. (**a**) Schematic for comparing exocytosis rates of axon-contacting vs. non-contacting oligodendrocytes that are co-cultured with active neurons. (**b**) Spatial distribution of VAMP3 exocytosis events (white dots) from contacting (left) and non-contacting (right) oligodendrocytes (blue) with axon overlay in magenta. (**c**) Exocytosis rates of VAMP3 in oligodendrocytes co-cultured with active neurons, stratified from Fig. 1d with the mean range of rates from isolated oligodendrocytes shown in the green overlay. p-value: unpaired, two-tailed t-test. (**d**) Exocytosis rates of VAMP2 and VAMP3 in isolated oligodendrocytes acutely treated with 0 mM (−) or 3 mM (+) glutamate for 30 min. p-values: one-way ANOVA with Tukey correction for multiple comparisons. (**e**) Exocytosis rates of VAMP3 in isolated oligodendrocytes treated with vehicle, NBQX, MK-801, and MTEP (30 min) prior to acute glutamate (3 mM) treatment. p-values: one-way ANOVA with Tukey correction for multiple comparisons.

### Exocytosis in mature oligodendrocytes drives activity-dependent myelin remodeling

Because oligodendrocytes rely on exocytosis to add new myelin membrane during sheath elongation in development^16^, we asked whether activity-gated exocytosis in mature oligodendrocytes drives myelin remodeling during adulthood. We used an optogenetic approach to induce myelin remodeling by activating callosal projection neurons in adult Thy1::channelrhodopsin 2 (ChR2) mice^27^. Chronic, unilateral optogenetic stimulation of layer V projection neurons in the premotor cortex has been shown to increase myelin thickness on axons projecting into the corpus callosum by 4 weeks post-stimulation^11,28^, providing a model with regionally and temporally defined parameters for tracking myelin changes (Fig. 3b). Increasing myelin sheath thickness through optogenetic stimulation requires BDNF-TrkB signaling between neurons and oligodendrocyte precursors, and is associated with generation of new oligodendrocytes^11^, but how stimulating neuronal activity affects sheath length and pre-existing myelin remains incompletely understood. To specifically label pre-existing sheaths that myelinate the light-stimulated neurons, we injected AAV expressing a membrane-bound fluorescent reporter under the oligodendrocyte-specific Mbp promoter^29^ (pMBP-mRuby-caax) into the corpus callosum ventral to the optical fiber (Fig. 3a). At 3-4 weeks post-stimulation, we harvested brain tissue to trace myelin sheaths on sparsely labeled oligodendrocytes ipsilateral to the stimulation site (Fig. 3c). Myelin sheaths in light-stimulated mice were 32% longer on average (47.1 to 62.0 μm) than in unstimulated mice (Fig. 3e-f), demonstrating that neuronal activity elongates pre-existing myelin sheaths in this paradigm.

**Fig. 3.**
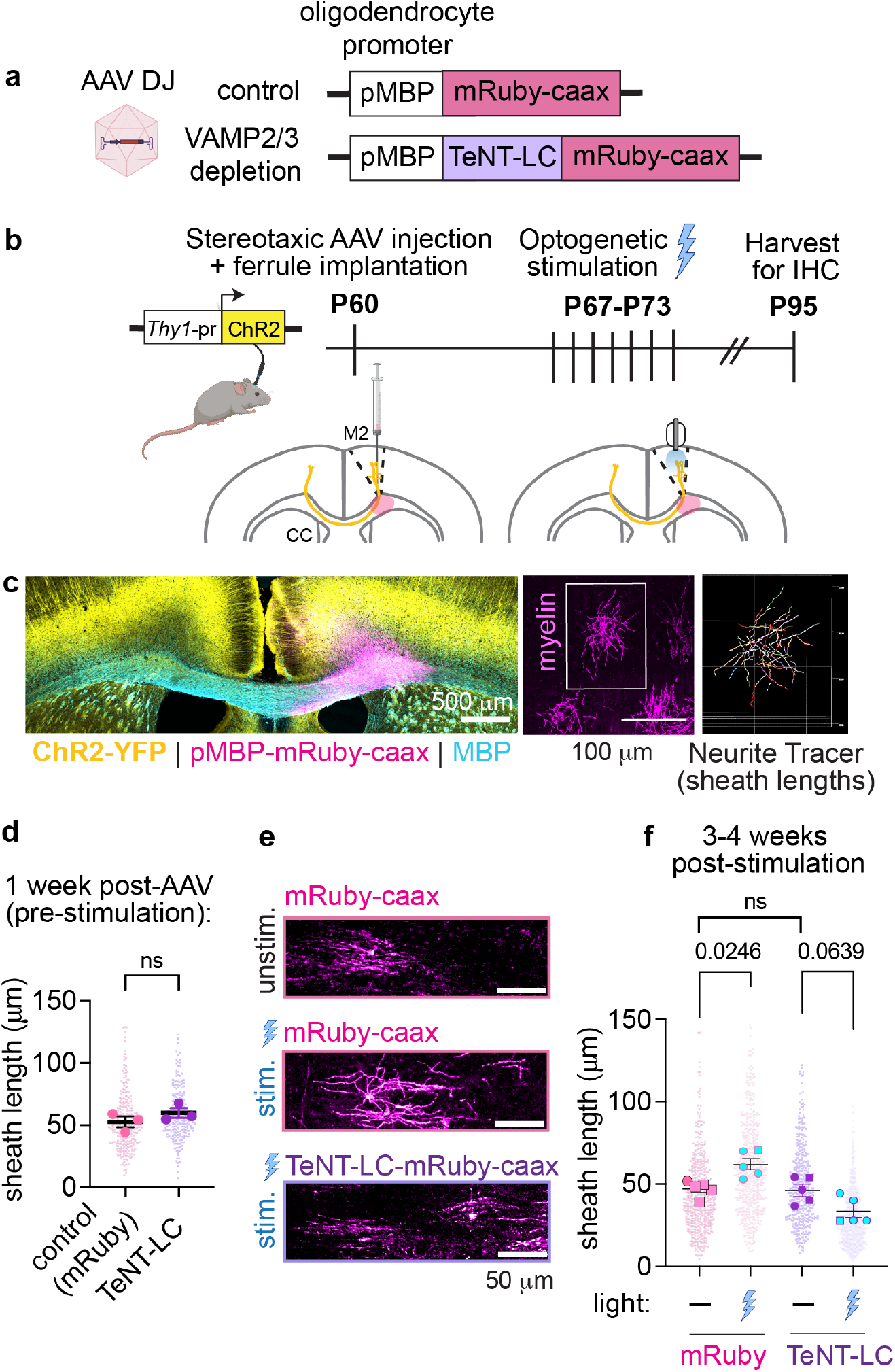
Oligodendrocyte exocytosis is necessary for activity-dependent myelin sheath elongation in the adult brain. (**a**) Adeno-associated viral (AAV-DJ) tools to label pre-existing oligodendrocytes in white matter **(b)** Experimental timeline for AAV-mediated labeling of pre-existing myelin in the corpus callosum (CC) followed by optogenetic stimulation (488-nm pulses at 20 Hz) of neurons in the pre-motor cortex (M2) transgenic mice. **(c)** Left: Confocal micrograph from a Thy1::ChR2 mouse with labeled oligodendrocytes in the corpus callosum stained by myelin basic protein (MBP). Middle: A single oligodendrocyte in the deep cortical layers/corpus callosum. Right: 3D reconstruction of labeled sheaths using Neurite Tracer. (**d**) Sheath lengths at 1 week post-AAV injection in unstimulated Thy1::ChR2 mice (P67, N=3 per group). p-value: unpaired, two-tailed t-test. (**e**) Labeled oligodendrocytes expressing the control or TeNT-LC-conjugated mRuby-caax protein from unstimulated or light-stimulated Thy1::ChR2 mice. (**f**) Sheath lengths at 3-4 weeks post-stimulation paradigm in *Thy1*::ChR2 mice (P95, N=5 per group). p-values: one-way ANOVA with Tukey correction for multiple comparisons. See Supplementary Table 2 for average sheath lengths ± SEM.

Does activity-dependent sheath elongation require exocytosis in pre-existing myelin? We replaced the control reporter with an AAV expressing tetanus toxin light chain (pMBP-TeNT-LC-mRuby-caax), which degrades VAMP2 and VAMP3 by cleaving between their cytosolic and transmembrane domains^30^ (Fig. 3a). Prior to stimulation, the average sheath length did not differ between control-and TeNT-labeled oligodendrocytes (Fig. 3d). However, optogenetic stimulation failed to increase—and instead trended toward decreasing—sheath length in TeNT-labeled oligodendrocytes (Fig. 3e-f), demonstrating that VAMP2/3-mediated exocytosis drives activity-dependent myelin remodeling.

### Remodeling of pre-existing myelin sheaths is required for motor learning

Finally, with VAMP2/3-mediated exocytosis as a genetic handle, we asked whether myelin remodeling is required for learning and/or memory. To degrade VAMP2/3 in pre-existing myelin during adulthood, we crossed mice with Cre-driven expression of botulinum toxin light chain (iBot)^31^ to mice with tamoxifen-inducible Cre in mature oligodendrocytes (*Plp*-CreERT)^32^ (Fig. 4a-b). At 6 weeks after tamoxifen injections, iBot expression was at least 88% specific to oligodendrocyte-lineage cells, with no loss of oligodendrocytes and widespread cleavage of VAMP2/3^33,34^ (Extended Data Fig. 3a-e). iBot; *Plp*-CreERT mice and littermate controls made similar numbers of reach attempts throughout the 10-day learning period, indicating no gross effect on motor ability or motivation (Extended Data Fig. 2a-b). However, iBot; *Plp*-CreERT mice had lower rates of successful reaches than littermate controls by the end of learning and maintained a lower performance during recall (Fig. 4e-f), revealing that remodeling of pre-existing myelin sheaths is required for motor learning in adulthood. In contrast, adult mice with VAMP2/3 degraded in oligodendrocyte precursors (iBot; *Pdgfra*-CreERT)^35^ achieved similar success rates as control littermates during learning (Fig. 4g) with no differences in reach attempts (Extended Data Fig. 2c-d). Tamoxifen-induced iBot expression in these mice was 86% specific to oligodendrocyte-lineage cells and impaired the generation of new oligodendrocytes post-training (Extended Data Fig. 4). Despite learning the task normally, iBot; *Pdgfra*-CreERT mice performed worse during recall (Fig. 4h), suggesting that new oligodendrogenesis may be necessary for longer-term motor skill maintenance independently of learning.

**Fig. 4.**
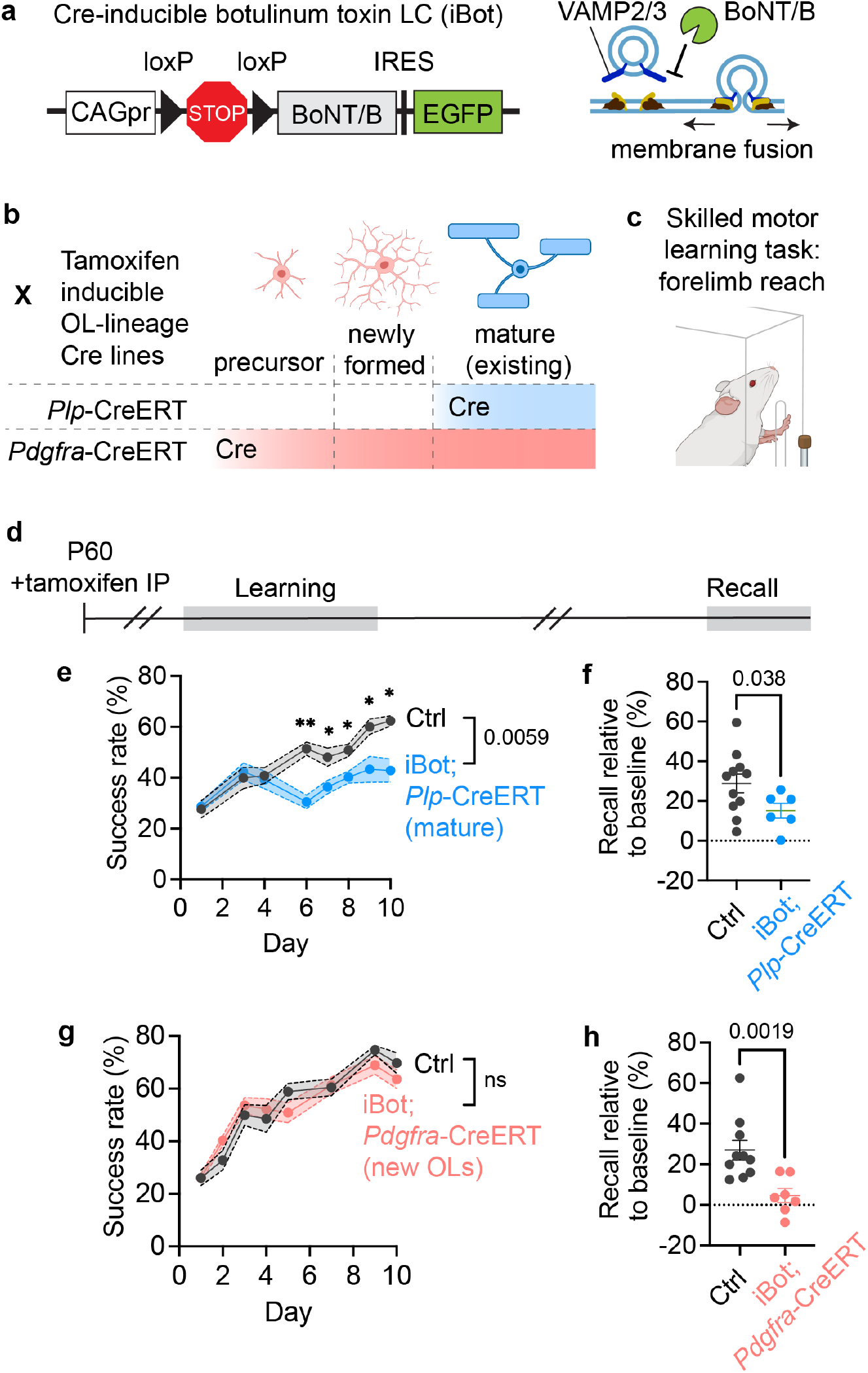
Exocytosis in existing myelin sheaths is required for proficient motor learning during adulthood. (**a**) Transgenic mouse for Cre-inducible blockade of exocytosis through botulinum toxin light chain B (BoNT/B) (**b**) Genetic crosses to obtain tamoxifen-inducible expression of iBot in existing (*Plp*-CreERT) or newly formed/precursor (*Pdgfra*-CreERT) oligodendrocytes (**c**) Single-pellet forelimb reach task as assay for motor skill learning. **(d)** Timeline for iBot induction and motor skill learning followed by recall. (**e**,**g**) Rate of successful pellet retrievals over training day. Ctrl N=11, iBot; *Plp*-CreERT N=6. Ctrl N=10, iBot; *Pdgfra*-CreERT N=7. p-values: mixed effects model with Tukey correction for multiple comparisons. (**f**,**h**) Change in rate of success from Day 0-2 of training to recall (3 weeks post-training). p-values: unpaired Welch’s t-test.

## Discussion

Myelin plasticity allows dynamic fine-tuning of neural circuit timing by inducing new oligodendrocyte generation or by remodeling pre-existing sheaths. Activity-dependent cues that induce new oligodendrocyte generation are becoming clear^11,36^, but mechanisms that couple neuronal activity to myelin remodeling and their causal contribution to learning remain to be discovered. Our work demonstrates that glutamatergic neuronal activity stimulates exocytosis in mature oligodendrocytes to drive remodeling of pre-existing myelin sheaths. Given our previous findings that exocytosis is essential for myelin membrane expansion^16^, we posit that activity-gated exocytosis directly adds new myelin membrane to adjust sheath morphology. We leveraged this pathway to block myelin remodeling in adult mice and found that exocytosis in pre-existing myelin is required for motor learning. Intriguingly, exocytosis in newly formed oligodendrocytes and their precursors is instead required for memory of the motor task, but not for learning. Altogether, our results uncover an activity-gated mechanism in pre-existing myelin that enables learning, presenting oligodendrocyte exocytosis as a driver of structural plasticity in the adult brain.

Do our measured rates of activity-gated exocytosis in oligodendrocytes align with the timeframe of myelin remodeling and motor learning? A typical oligodendrocyte in the mouse corpus callosum can have up to 30 sheaths^37^ with 10 wraps^38^ each around axons of 1 μm in diameter. We measured that optogenetic stimulation of neurons increased the average myelin sheath length by 32% (from 47 μm to 62 μm, Fig. 3e-f) over 28 days. If every single sheath layer increased by 32% in length (a maximal assumption), a mature oligodendrocyte would need ~14,500 μm^2^ of new membrane area, amounting to 460,000-1,800,000 new vesicles (with spherical diameters of 50-100 nm). At the measured rate of 50 vesicles/min (the activity-gated increase in VAMP3 exocytosis, Fig. 1d), incorporating this range of vesicles would take 6-25 days. This timeframe aligns with the protracted process of remodeling existing sheaths induced by optogenetic stimulation, by sensory experience^5^, and by motor learning^4^. More interestingly, this timeframe aligns with the emergence of learning deficits in mice with disrupted exocytosis in existing myelin (Fig. 4d), suggesting that activity-gated exocytosis in oligodendrocytes modulates neural circuits on behaviorally-relevant timescales.

On the other hand, how does neuronal activity, occurring at millisecond timescales, elicit protracted membrane dynamics in myelin? Our live-cell imaging of oligodendrocytes reveals that VAMP3 exocytosis is calcium-dependent and stimulated by glutamate through metabotropic glutamate receptor 5 (mGluR5). In rodent brain tissue, evoked action potentials trigger axonal release of glutamate in white matter tracts^39,40^, providing a localized source of glutamate for existing myelin. In larval zebrafish, mGluR5 was recently identified as the key glutamate receptor in myelin sheaths that mediates activity-induced calcium transients and sheath elongation^21^. Metabotropic signaling via mGluR5 offers a path to convert millisecond changes in neuronal activity to prolonged calcium transients that develop over several seconds to hours^41^. Adult mice with attenuated calcium signaling in oligodendrocytes exhibit aberrant myelin morphology^24^, and our preliminary data indicate that calcium signaling in mature oligodendrocytes is required for proficient motor learning^42^. How calcium signaling sculpts myelin sheath morphology remains an active area of research^22,42^. Our findings present a missing link between glutamate-induced calcium transients and myelin membrane dynamics by identifying VAMP3 as the downstream driver of membrane addition to elongate myelin sheaths.

Importantly, our data lay the groundwork for mechanistic studies into the rules and principles of myelin remodeling across different circuits and adaptive behaviors. Although we have identified glutamate as a trigger for exocytosis and sheath elongation, myelin remodeling also occurs on parvalbumin interneurons upon changes in sensory experience^5^. Whether non-glutamatergic neurons modulate exocytosis in oligodendrocytes remains an open question. In addition to sheath elongation, neuronal activity can also induce shortening of existing sheaths, which lengthens Nodes of Ranvier to modulate conduction velocity^4^. Sheath shortening in Layers I and II/III of the motor cortex correlated with learning performance in the single-pellet forelimb reach task^4^. Determining whether oligodendrocyte exocytosis contributes to myelin changes in the motor cortex or mediates sheath elongation in a distinct region will reveal how different mechanisms of myelin remodeling are coordinated across brain regions. Intriguingly, we found that sheaths with blocked exocytosis trended toward shortening under stimulation rather than merely failing to grow (Fig. 3f), suggesting that neuronal activity “unlocks” myelin sheath plasticity more broadly, with exocytosis determining whether that plasticity resolves as growth. Because a single oligodendrocyte myelinates multiple, often functionally distinct axons, gating exocytosis locally at individual sheaths, rather than uniformly across the cell, would let one oligodendrocyte tune each individual sheath to its own axon’s activity.

Lastly, how do myelin remodeling and new oligodendrocyte generation coordinate to support learning and memory? Our genetic dissection of disrupting exocytosis in mature vs. precursors of oligodendrocytes suggests dissociable roles in motor learning and memory maintenance. Disrupting exocytosis in oligodendrocyte precursors (iBot; *Pdgfra*-CreERT) reduced new oligodendrocyte generation (consistent with the role of exocytosis in promoting oligodendrocyte differentiation^43,44^) and impaired motor memory without affecting learning. Although new oligodendrocyte generation is required for certain motor learning tasks^1^, a recent study also found that new oligodendrocyte generation is not required for learning the single-pellet forelimb reach task^45^. Instead, genetically ablating the ability to form new oligodendrocytes reduced memory of the skill^45^, consistent with our findings using disruption of exocytosis. Longitudinal imaging also showed that new oligodendrocyte generation is most abundant after the cessation of learning^46^ when new sheath addition occurs^4^ to presumably consolidate changes to neural circuits.

Additionally, exocytosis in oligodendrocyte precursors may be important to stabilize pre-synapses on axons in an activity-dependent manner as an orthogonal mechanism to sculpt neural circuitry^47^. Given the distinct roles of exocytosis in oligodendrocytes and their precursors, exocytosis can be used as a genetic handle to compare the relative contributions of myelin remodeling, new oligodendrocyte generation, and OPC-axon synapses within the same adaptive behavior.

Our data identify a cellular mechanism that directly couples neuronal activity to myelin membrane addition in pre-existing sheaths, opening new avenues to modulate myelin remodeling in specific circuits or disease contexts. We anticipate that inhibiting oligodendrocyte exocytosis will provide a strategy to reveal the functional role of myelin dynamics in axonal health and circuit activity, while stimulating this pathway may enhance myelin repair in injury or disease.

## Acknowledgements

We thank Dr. Erin Gibson for helpful insight on the experimental design of the optogenetic stimulation for myelin remodeling; Drs. Frank Pfrieger and Shane Hentges for the iBot mouse line; Dr. Stephanie Gupton for the VAMP-pHluorin plasmids; and Dr. Brigitte Dorner for the cleaved VAMP2/3 antibody (VAMP/B/151). Images and diagrams were created using BioRender.com. We also thank the Stanford Neuroscience Gene Vector and Virus Core for producing the adeno-associated viruses (pMBP-mRuby-caax and pMBP-TeNT-LC-mRuby-caax) used in this study.

This project was supported by the National Institutes of Health K99NS136783 (M.L.) and R01NS134829 (J.B.Z. and E.G.H.), a Stanford Alzheimer’s Disease Research Center Research Education Component (ADRC REC) grant (M.L.), and the McKnight Endowment Fund for Neuroscience Neurobiology of Brain Disorders Award (J.B.Z. and E.G.H.).

This paper was typeset with the bioRxiv word template by @Chrelli: www.github.com/chrelli/bioRxiv-word-template.

## Author Contributions

M.L. and J.B.Z. conceived of the project. M.L. designed, performed, and analyzed all experiments with the following exceptions. O.D. performed and analyzed the live-cell imaging of isolated oligodendrocytes treated with glutamate and glutamate receptor antagonists in Fig. 2d-e. O.D. also harvested brains, sectioned and stained tissue slices for immunohistochemistry, and acquired microscopy images of transgenic mouse brains for Extended Data Fig. 3-4. A.C.G, B.Y. and M.M. designed and trained M.L. on experiments using optogenetic stimulation in Fig. 3. L.D. acquired and analyzed biological replicates of VAMP2 and VAMP3 exocytosis in Fig. 1c-d. M.I. designed and created the construct for genetically encoded calcium chelation in oligodendrocytes used in Fig. 1e-g. K.H.S and E.G.H. provided experimental rationale for testing transgenic mice on the forelimb reach learning assay based on prior results, training on the assay, and input on the design and analysis of the behavioral assay. M.L. and J.B.Z. wrote the manuscript. All authors reviewed and gave feedback on the manuscript.

## Materials and Methods

### Mouse experiments

All animal procedures were approved by Stanford University’s Administrative Panel on Laboratory Animal Care (APLAC) and followed the National Institutes of Health guidelines. Mice were group-housed under standard 12:12 light-dark cycles with free access to food and water, and disposable bedding in plastic cages. All mice were regularly monitored by veterinary and animal care staff and were not involved in prior procedures or testing. Sprague-Dawley rats and C57BL/6 mice were ordered from Charles River Laboratories. Both male and female mice were studied for all in vivo experiments.

The transgenic iBot mice line, which contains the floxed-STOP site upstream of the Clostridium botulinum neurotoxin light chain followed by an IRES-GFP, was kindly gifted by Dr. Frank Pfrieger (University of Strasbourg) and Dr. Shane Hentges (Colorado State University). The transgenic tamoxifen inducible Cre lines were obtained from Jax: B6.Cg-Tg(Plp1-cre/ERT)3Pop/J (Strain #:005975) and B6N.Cg-Tg(Pdgfra-cre/ERT)467Dbe/J (Strain #:018280).

All mice were injected intraperitoneally with 100 mg/kg tamoxifen (Sigma T5648) dissolved in corn oil for 5 consecutive days starting at post-natal day 60 (P60) of age and trained 6 weeks later. Males and females were pooled for analysis because no sexual dimorphism for myelination defects was observed after the conditional expression of iBot.

### Preparation of re-aggregated retinal ganglion cells (RGCs)

Our protocol for myelinating co-cultures with CNS-derived axons was adapted from previous publications^16,17^ with minor modifications. Dissection and dissociation of retina, immunopanning for retinal ganglion cells (RGCs) with anti-Thy 1.1, and media composition with growth factor supplementation were performed as previously published.

To create re-aggregates, we seeded freshly harvested RGCs into UV-sterilized PCR tubes (Fisherbrand 14-230-215) at a density of 10,000 cells/100 μl of media. After 24 h of recovery, we half-fed each tube by exchanging 50 μl of media with freshly supplemented growth factors. After 48 h of recovery, we transferred 2x 100 μl cell suspension containing “re-aggregated” clumps of RGCs from each tube onto one well of a PDL-borate-coated 8-well iBidi slide (iBidi Cat. 80826) to set up for live-cell imaging. Each well was half-fed with fresh growth factors every 72–96 h. After 10–14 days, the RGC re-aggregates formed dense beds of radially protruding axons.

### Co-culturing of transfected oligodendrocytes with neurons (RGCs)

Primary oligodendrocyte precursors were purified by immunopanning from P5-P7 Sprague Dawley rat as previously described (Dugas and Emery, 2013; Emery and Dugas, 2013). Oligodendrocyte precursors were typically seeded at a density of 150,000-250,000 cells/10-cm dish and allowed to recover for 4 days in culture before lifting cells via trypsinization and distributing for transfection, proliferation, or differentiation assays. All plasticware for culturing oligodendrocyte precursors were coated with 0.01 mg/ml poly-D-lysine hydrobromide (PDL, Sigma P6407) resuspended in water. All glass coverslips for culturing oligodendrocyte precursors were coated with 0.01 mg/ml PDL, which was first resuspended at 100x in 150 mM boric acid pH 8.4 (PDL-borate).

To proliferate primary oligodendrocyte precursors, cells were cultured in serum-free defined media (DMEM-SATO base medium) supplemented with 4.2 μg/ml forskolin (Sigma-Aldrich, Cat#F6886), 10 ng/ml PDGF (Peprotech, Cat#100-13A), 10 ng/ml CNTF (Peprotech, Cat#450-02), and 1 ng/ml neurotrophin-3 (NT-3; Peprotech, Cat#450-03) at 37°C with 10% CO2. To induce differentiation, cells were switched to DMEM-SATO base media containing 4.2 μg/ml forskolin (Sigma-Aldrich, Cat#F6886), 10 ng/ml CNTF (Peprotech, Cat#450-02), 40 ng/ml thyroid hormone (T3; Sigma-Aldrich, Cat#T6397), and 1x NS21-MAX (R&D Systems AR008).

Proliferating rat oligodendrocyte precursors were lifted from tissue culture dishes, and centrifuged at 90xg for 10 min. 250,000 oligodendrocyte precursors were gently resuspended into 20 ul of nucleofector solution (Lonza P3 Primary Cell 4D-Nucleofector V4XP-3032) with 300 ng of VAMP-pHluorin and 300 ng of mRuby-caax plasmids. Cells were then electroporated in a Lonza 4D-Nucleofector X Unit (AAF-1003X) assembled with a 4D-Nucleofector Core Unit (AAF-1002B) with pulse code DC-218. Electroporated cells rested for 10 min at RT before resuspension in antibiotic-free DMEM-SATO base media supplemented with differentiation factors. Each batch of 250,000 cells was distributed into 4 wells of an 8-well iBidi slide containing media alone or RGC reaggregates.

### Live-cell imaging of co-cultures and isolated oligodendrocytes

Time-lapse imaging of exocytic events was performed on a Zeiss Axio Observer Z1 inverted microscope equipped with a Zeiss Axiocam 506 monochrome 6-megapixel camera, a stage top incubator (Okolab, H301-K-Frame) set to 37 °C, and a digital gas blender (Okolab, CO2-UNIT-3L) set to 10% CO2 during image acquisition. Samples were imaged with a Plan-Apo 63×/1.40 Oil objective using widefield epifluorescence with a 12 V halogen lamp. Then time-lapse sequences of exocytotic events were captured at 250 ms/frame for 1 min with Zen Blue software. Images were viewed using Fiji/ImageJ software.

### Acute glutamate treatment of primary oligodendrocytes

The imaging medium consisted of FluoroBrite DMEM supplemented with SATO, 2 mM MgCl_2_, forskolin, CNTF and T3 at final concentrations of 1:1000, sterile-filtered and prepared fresh on the day of imaging. A 100 mM L-glutamate stock was likewise prepared fresh immediately before each experiment: 44.2 mg L-glutamic acid (MW 147.13) was dissolved in 2.5 mL imaging medium with 150 μl of 1 N NaOH, warmed to 37 °C to bring the free acid into solution, adjusted to pH ~7.0, and brought to a final volume of 3.0 mL with imaging medium. This stock was diluted into 1.5 mL aliquots of imaging medium to yield working concentrations alongside a vehicle-only aliquot. Immediately before imaging, the medium was exchanged for 350 μl of the appropriate glutamate (for a final concentration of 3 mM) or vehicle solution as the iBidi chamber was transferred to the microscope. Slides were equilibrated on the stage in a 37 °C, 10% CO_2_ environmental chamber for 15 min before acquisition to allow thermal stabilization and to permit the glutamate response to reach a steady rate. Each well was imaged within 30 min of glutamate addition to limit variability in exposure time across fields.

### Pharmacological inhibition of the glutamate response in primary oligodendrocytes

To determine which glutamate receptor mediates the stimulation of exocytosis, DIV5 cells were pre-incubated with receptor antagonists for 30 min before glutamate was applied and imaged 15 min after glutamate addition, giving a total of 45 min on the stage for thermal stabilization. Antagonists were diluted in imaging medium and applied by medium exchange at the following final concentrations: MTEP (Tocris 2921; 80 μM), NBQX (Tocris 1044; 50 μM), and MK-801 (Tocris 0924; 200 μM). After the 30 min pre-incubation, the medium was exchanged a second time for imaging medium containing the same concentration of antagonist together with L-glutamate (Sigma-Aldrich G1251) at 3 mM, freshly prepared as described above. Control wells underwent an identical two-step medium exchange with antagonist- and glutamate-free imaging medium and the same 30 min + 15 min waiting period, so that all conditions were matched for handling, medium exchange, and total time on the stage. This control accounts for any exocytotic response evoked by mechanical perturbation or medium change alone. Each antagonist condition and its paired control were imaged from the same 8-well μ-slide and the same culture preparation on a given day, and conditions were acquired in randomized order to control for drift in cell health over the imaging session.

### Analysis of VAMP2/3-mediated exocytotic events in primary oligodendrocyte cultures

Exocytotic events were quantified using IVEA (Intelligent Vesicle Event Analysis)^48^, a deep learning–based pipeline that identifies exocytotic events as non-motile, Gaussian-shaped puncta undergoing transient fluorescence increases against the local background of the pHluorin time-lapse. Time-lapse recordings of VAMP2/3-pHluorin–expressing oligodendrocytes at DIV5 were imported as single-plane videos acquired at 4 Hz with a pixel size of 72.0635 nm, and processed with the Random Burst Event module using the GranuVision3 convolutional network. Candidate events were nominated within a vision radius of 12 pixels and tracked with a search radius of 3 pixels, with no temporal max pooling (pooling factor = 1) so that every frame was evaluated independently. Event nomination used a peak prominence of 10, a variation time window of 2 frames, an event spread of 0, and a nomination sensitivity of 1, with Gaussian blur applied to suppress shot noise; candidates were retained as exocytotic events only when classified by the network with a confidence ≥0.85. Detection was configured to include events lacking a diffusive fluorescence cloud (class 1) and latent vesicle fusion events (class 2), as well as small/low-amplitude events, to capture the full range of VAMP2/3-dependent fusion behaviors in oligodendrocytes; because the last of these settings is permissive to noise, all nominated events were subsequently inspected manually and false positives arising from focal drift, membrane ruffling, or debris were discarded. For each retained event, fluorescence was measured over a window of 20 frames before to 150 frames after the nominated fusion frame, from which event amplitude and decay kinetics were extracted. Event frequencies were expressed as events per cell per minute.

### Stereotaxic surgery and optogenetic stimulation

Serotype AAV-DJ was prepared by the Stanford Neuroscience Gene Vector and Virus Core, and stored at−80 °C until use.

At P60, mice were anesthetized with 1–4% isoflurane and secured in a stereotactic frame. Following aseptic preparation of the scalp, a midline incision exposed the skull, and a 0.5-mm burr was used to thin the bone over the target region (0.5 mm lateral to midline, 1.0 mm anterior to bregma). AAV encoding either pMBP-mRuby-caax (control) or pMBP-TeNT-LC-mRuby-caax (1 μl of 2.5 E+12 vg/ml) was injected into the corpus callosum (−1.5 mm ventral). A fiber optic ferrule (Doric Lenses) was positioned above the right premotor cortex (M2) at coordinates 0.5 mm lateral to midline, 1.0 mm anterior to bregma, and 0.6 mm ventral to the skull surface, then fixed in place with dental cement.

Starting on P67, mice were tethered by a monofiber patch cord to a 473-nm diode-pumped solid-state laser and allowed to move freely during stimulation. Blue light was delivered at 20 Hz (50-ms cycles; 25-ms on/25-ms off) at an intensity of ~1 mW measured at the fiber tip, matching the parameters reported by Gibson et al.^28^ to activate this circuit. Each daily session consisted of repeating 30-s stimulation epochs separated by 2-min rest intervals, totaling 10 min of cumulative light exposure per day, over 7 consecutive days (P67–P74). Mice were euthanized at P95 for tissue collection, 3 weeks after the final stimulation session.

A parallel cohort of control mice received the same surgical procedure and ferrule implantation but was never exposed to blue light stimulation.

### Immunohistochemistry of mouse brain tissue

For histological analysis, mice were anesthetized by injecting 2.5% avertin (100 ul/10 g). The animals were then transcardially perfused with PBS and then with 4% paraformaldehyde (PFA) (diluted into 1xPBS from 16% PFA, Electron Microscopy Sciences), followed by post-fixation with 4% PFA for immunohistochemistry. Then brains were washed in PBS, cryoprotected in 30% sucrose for 48-72 hrs, and embedded in OCT; blocks were stored at −80°C and transferred to −20°C the day before sectioning to equilibrate. A fiducial nick was made with a 27G needle in the hemisphere ipsilateral to the reaching forelimb to preserve ipsi/contralateral orientation throughout processing. Blocks were sectioned on a cryostat (CM3050 S, Leica Microsystems) into 50 μm free-floating coronal sections, which were collected in PBS and stored at 4°C for up to 24 h before staining. For ASPA labeling, heat-mediated antigen retrieval was performed before staining by incubating free-floating sections in 1× citrate buffer, pH 6.0 (C9999-100ML, diluted 1:10 in water) for 30 min at 60°C in a rotating heat block, followed by 3 × 5 min washes in PBS with agitation. Sections were then blocked in 3% normal donkey serum (ab7475) with 0.3% Triton X-100 in PBS for 1 h at RT with agitation and washed 3 × 10 min in PBS. Primary antibodies were diluted in 1% normal donkey serum with 0.3% Triton X-100 in PBS (400 μl per well of a 24-well plate) and incubated at 4°C for 72 h. To assess the penetrance and cell-type specificity of iBot expression, sections were labelled with chicken anti-GFP (1:500), goat anti-Olig2 (1:250) and rabbit anti-ASPA (1:500). Primary antibody was washed out with 3 × 10 min washes in 2 ml PBS per well at RT with agitation, and donkey secondary antibodies diluted in 1% normal donkey serum with 0.3% Triton X-100 in PBS were applied overnight at 4 °C: anti-chicken Alexa Fluor 488 (1:1000), anti-rabbit Alexa Fluor 568 (1:500), and anti-goat or anti-rat Alexa Fluor 647 (1:500). After 3 × 10 min PBS washes, sections were mounted onto Superfrost Plus (VWR) slides by floating them from a PBS-filled dish and arranging them with a #0 paintbrush, coverslipped in Vectashield with DAPI (Vector, H-1200), and sealed with nail polish. Antibody sources and catalogue numbers are specified in the Antibodies section.

### Antibodies and Staining Reagents

Primary antibodies used in this study were as follows: rat-anti-MBP (Abcam ab7349; 1:1000), goat-anti-Olig2 (R&D Systems AF2418; 1:250), chicken-anti-GFP (Abcam ab13970; 1:500), Rabbit-anti-Aspartoacylase (GeneTex GTX113389, 1:500). The antibody that recognizes the C-terminal fragment of cleaved VAMP2/3 (VAMP/B/151) was a generous gift from Prof. Brigitte G. Dorner (Robert Koch Institute, Berlin, Germany).

Secondary antibodies used in this study were as follows at a 1:1000 dilution for both tissue and primary cell immunofluorescence: donkey anti-chicken 488 (Biotium 20166), donkey anti-rabbit 568 (Thermo Scientific A10042), donkey anti-goat 647 (Biotium 20048), donkey anti-rat (Thermo Scientific A48272).

### Quantification of myelin sheath length

Sheath length was measured in Fiji/ImageJ^49^ using the SNT plugin^50^, by manually tracing the mRuby signal; tracing was performed through the z-stack so that sheaths coursing obliquely through the tissue were measured in three dimensions rather than in projection, and path lengths were calibrated to the voxel dimensions of the acquisition. All tracing was performed blinded to genotype. 50 sheaths were measured per animal. Statistics were performed on biological replicates (n = 5 animals per group).

### Forelimb reach training

Prior to training, mice were weighed and given a single 20-min session in the training box to acclimate to the apparatus, followed by a 24-h period of food restriction. Pellets were presented through a narrow slot in the box wall, positioned on a shelf offset 1 cm forward and 1 mm to the right of the opening. Training then proceeded once daily for 10 days; each session ended after 20 min. All mice were trained to reach with their left forepaw. A trial was scored as successful if the mouse grasped the pellet and brought it back through the slot into the box. Food restriction was maintained across the training period to sustain motivation, with daily weight checks to confirm that no animal lost more than 10% of body weight. Mice that resorted to licking for pellet retrieval over the first two training days were excluded from analysis. All animals were trained blinded to genotype by the same experimenter to avoid confounds from experimenter-specific handling effects.

Recall was assessed 3 weeks after the last day of training, using three additional sessions run under the same 20-min cutoff and scored with the same success/error criteria as during training.

## Data Analysis and Statistics

All analysis was conducted blinded to the genotype. Data analysis and statistics were done using GraphPad Prism 11 software. Descriptive statistics (mean, standard error of mean, and N) were reported in Figure Legends or Supplementary Tables. For cellular assays, the biological replicate represented the mean of technical replicates from one animal.

## Data and materials availability

The data that support the findings of this study and step-by-step protocols are available from the corresponding or first authors upon request. Correspondence and requests for all other materials should be addressed to M.L or J.B.Z.

**Extended Data Fig. 1.**
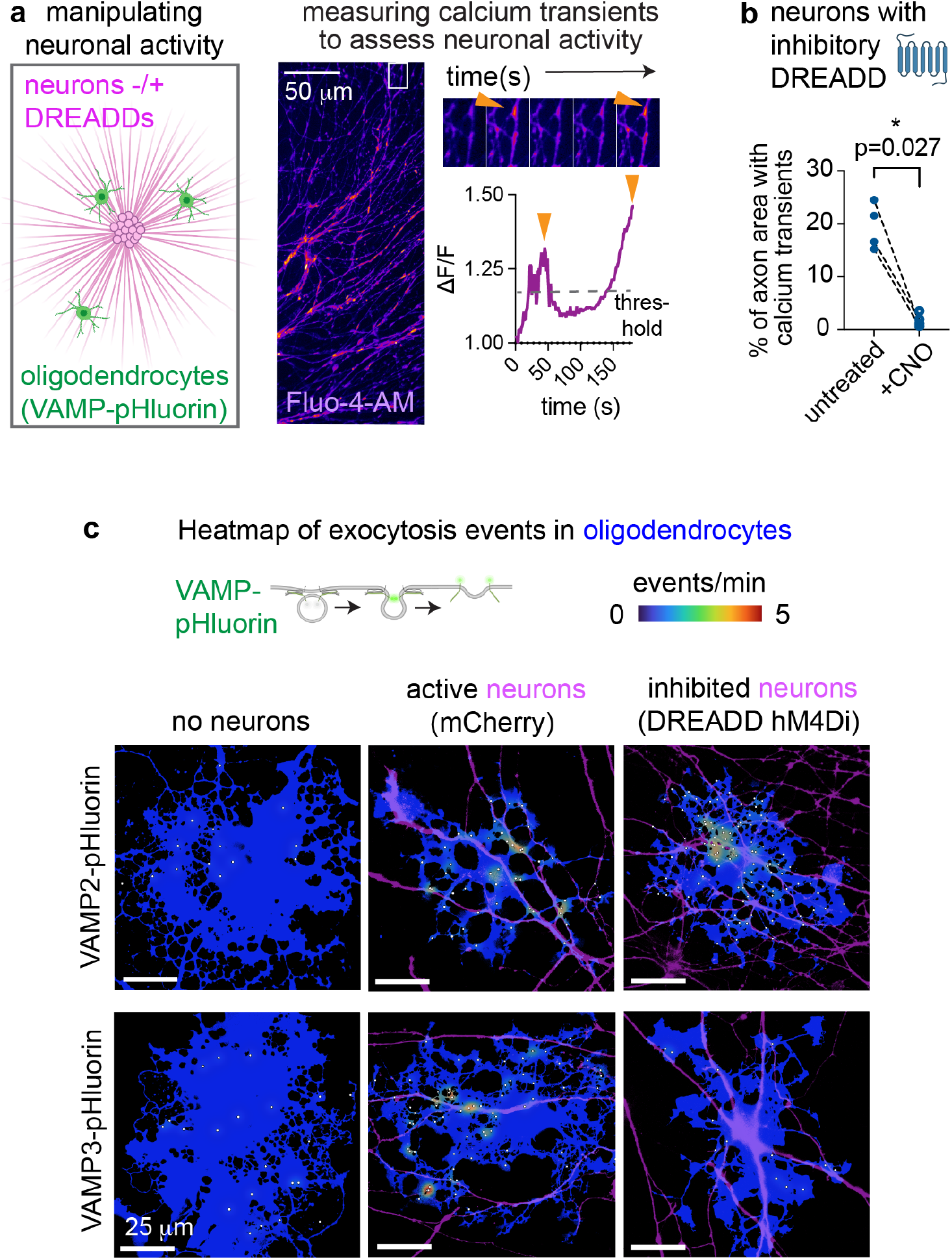
Spatial distribution of exocytosis events on oligodendrocytes co-cultured with neurons. (**a**) Time-lapse imaging of the calcium indicator Fluo-4-AM (500 nM) on retinal ganglion cell reaggregates. Example normalized fluorescence trace shows the thresholding used to bin positive calcium events. (**b**) Calcium transients before and after clozapine-N-oxide (CNO) treatment of DREADD-Gi infected neurons. p-value: two-tailed, paired t-test. (**c**) Heatmaps of VAMP2 (top) and VAMP3 (bottom) exocytosis events in oligodendrocytes (blue) on axon overlays (magenta).

**Extended Data Fig. 2.**
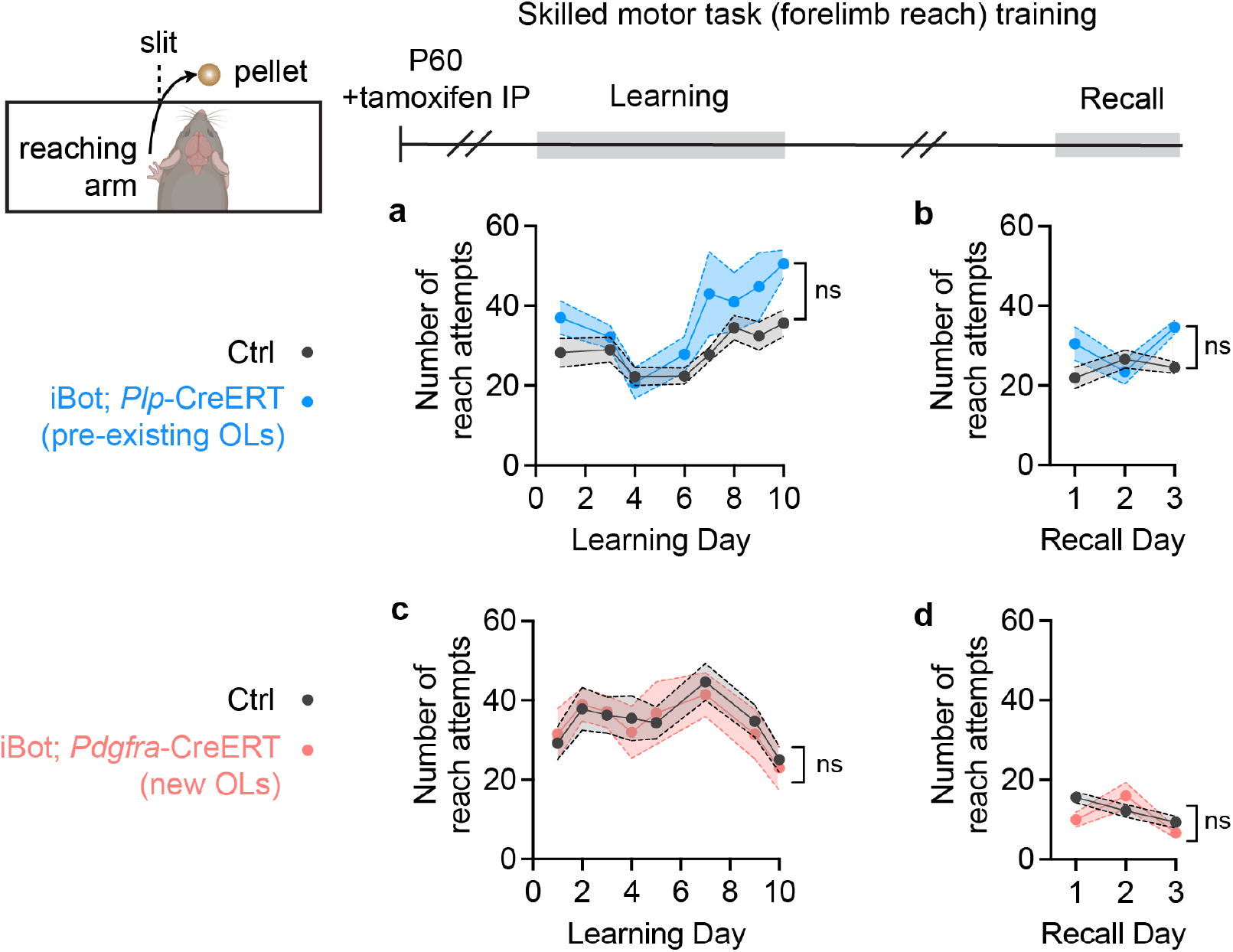
Depletion of VAMP2/3 does not affect reach attempts during learning or recall. (**a**,**c**) Number of reach attempts over training day of the single-pellet forelimb reach task. (**b**,**d**) Number of reach attempts over the recall period. Statistical analysis: mixed effects model with Tukey correction for multiple comparisons (ns = not significant).

**Extended Data Fig. 3.**
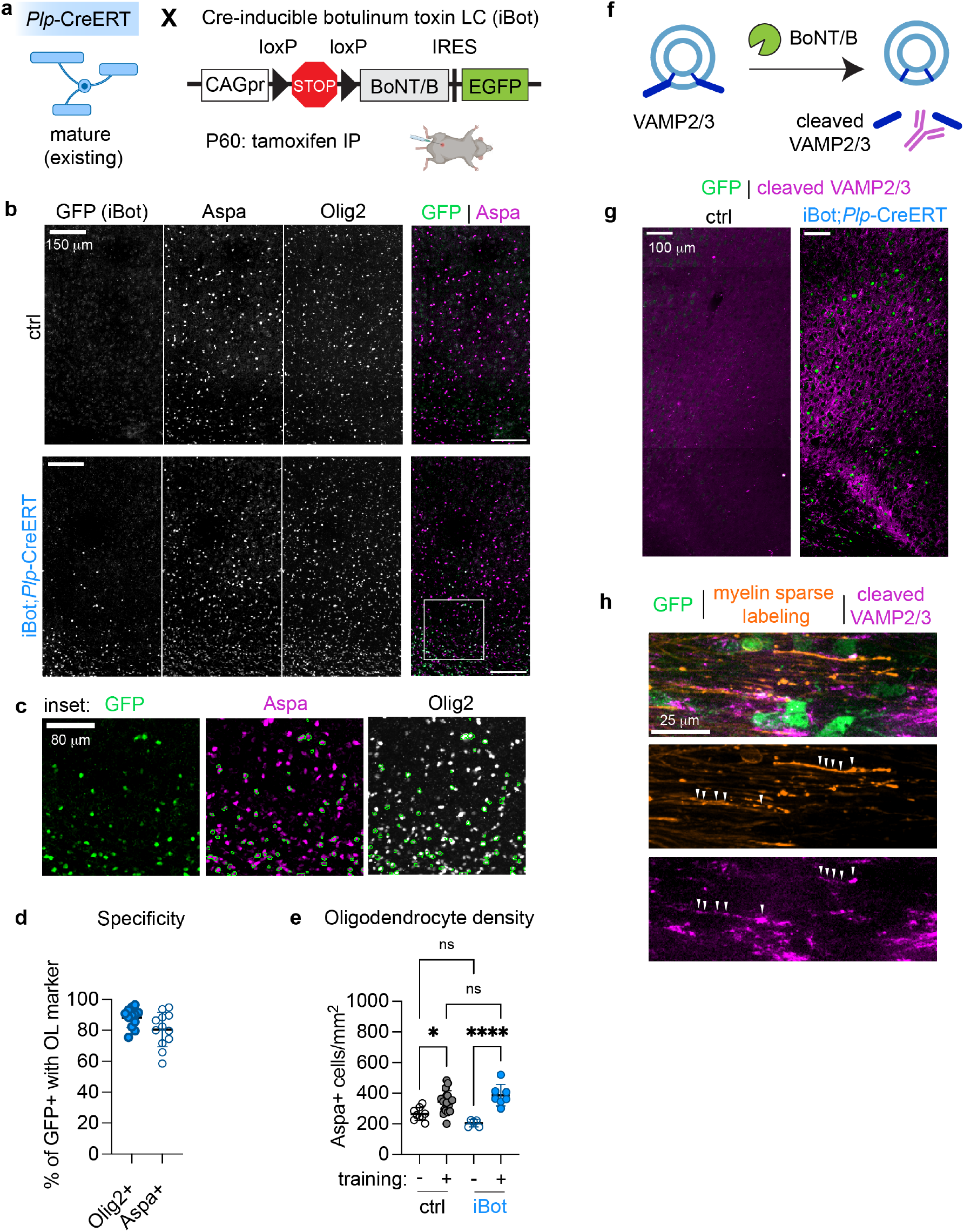
Disrupting VAMP2/3-mediated exocytosis in mature oligodendrocytes does not affect learning-induced changes in oligodendrocyte density. (**a**) Genetic cross for tamoxifen-inducible blockade of exocytosis in pre-existing myelin. (**b**) Stained coronal sections of motor cortex, where Aspa marks mature oligodendrocytes and Olig2 marks all oligodendrocyte-lineage cells. (**c**) Zoomed inset of GFP-Aspa merge in (b) with outlines of GFP+ cells. (**d**) Quantification of specificity of iBot expression marked by GFP. (**e**) Quantification of oligodendrocyte density (Aspa+) in motor cortex from trained and untrained ctrl and iBot;Plp-CreERT mice. p-values: one-way ANOVA with Tukey correction for multiple comparisons. (**f**) Schematic of botulinum neurotoxin B (BoNT/B)-mediated cleavage of VAMP2/3, generating a neoepitope recognized by the cleaved VAMP2/3 antibody. (**g**) Stained coronal sections from ctrl and iBot;Plp-CreERT mice showing GFP (iBot reporter) and cleaved VAMP2/3 immuno-reactivity. (**h**) Zoomed inset showing GFP, sparse myelin labeling, and cleaved VAMP2/3 in a single labeled oligodendrocyte; arrowheads indicate colocalization of cleaved VAMP2/3 with myelin sheaths.

**Extended Data Fig. 4.**
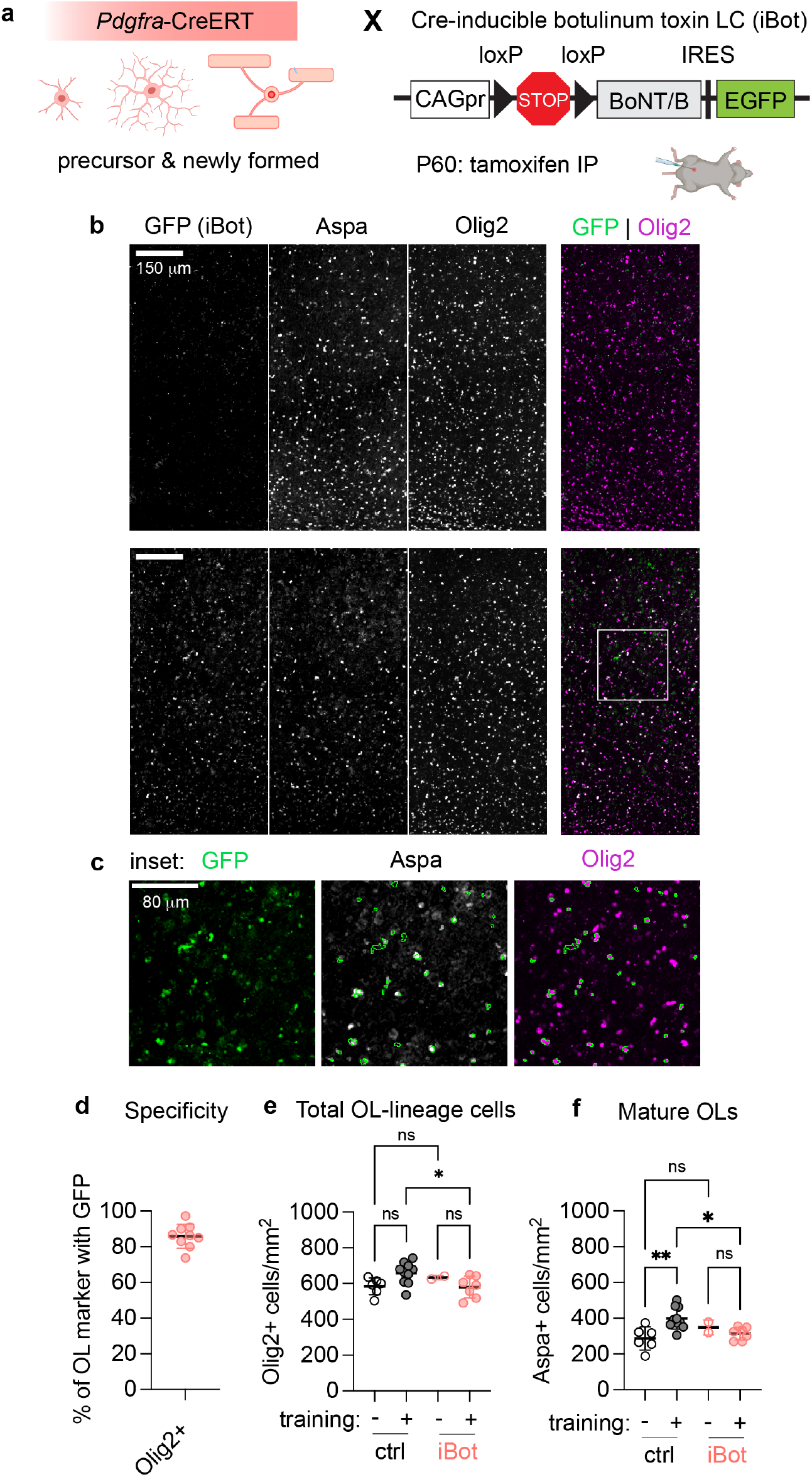
VAMP2/3-mediated exocytosis in oligodendrocyte precursors is required to increase oligodendrocyte density post-learning. (**a**) Genetic cross for tamoxifen-inducible blockade of exocytosis in newly formed myelin and oligodendrocyte precursors. (**b**) Stained coronal sections of motor cortex, where Aspa marks mature oligodendrocytes and Olig2 marks all oligodendrocyte-lineage cells. (**c**) Zoomed inset of GFP-Aspa merge in (b) with outlines of GFP+ cells. (**d**) Quantification of specificity of iBot expression marked by GFP. (**e**) Quantification of total oligodendrocyte-lineage cell density (Olig2+ cells/mm^2^) in motor cortex from trained and untrained ctrl and iBot;Pdgfra-CreERT mice. p-values: one-way ANOVA with Tukey correction for multiple comparisons. (**f**) Quantification of mature oligodendrocyte density (Aspa+ cells/mm^2^) in motor cortex from trained and untrained ctrl and iBot;Pdgfra-CreERT mice, showing reduced training-induced oligodendrogenesis in iBot;Pdgfra-CreERT mice. p-values: one-way ANOVA with Tukey correction for multiple comparisons.

**Supplementary Table 1:** Average exocytosis rates in oligodendrocytes co-cultured with neurons (RGC)

| Exocytosis in oligodendrocytes co-cultured with: |  | Vamp2 rates |  |  | Vamp3 rates |  |  |
| --- | --- | --- | --- | --- | --- | --- | --- |
|  |  | no neurons | active | inhibited | no neurons | active | inhibited |
| Reference Fig. 1c-d | Mean ± SEM | 25.4 ± 6.1 | 43.3 ± 3.9 | 39.4 ± 2.1 | 28.1 ± 5.7 | 79.3 ± 12.4 | 33.4 ± 8.12 |

| Exocytosis in oligodendrocytes (-/+ Ca<sup>2+</sup> chelator) co-cultured with active neurons: |  | Vamp2 rates |  | Vamp3 rates |  |
| --- | --- | --- | --- | --- | --- |
|  |  | control | SpiCee | control | SpiCee |
| Reference Fig. 1f-g | Mean ± SEM | 40.8 ± 1.3 | 39.6 ± 1.0 | 82.7 ± 7.5 | 31.0 ± 5.1 |

**Supplementary Table 2:** Average myelin sheath changes on optogenetically stimulated axons in the corpus callosum.

| Reference Fig. 3f | Sheath length (μm) of myelin labeled with: | mRuby (control) | | TeNT-LC | |
| --- | --- | --- | --- | --- | --- |
|  | Light stimulation | - | + | - | + |
| | Mean ± SEM | 47.1 ± 2.1 | 62.0 ± 3.6 | 46.2 ± 3.5 | 33.6 ± 3.6 |
|  | N (biological replicates) | 5 | 5 | 5 | 5 |

